# Structure-based drug repositioning explains ibrutinib as VEGFR2 inhibitor

**DOI:** 10.1101/678896

**Authors:** Melissa F. Adasme, Daniele Parisi, Kristien Van Belle, Sebastian Salentin, V. Joachim Haupt, Gary S. Jennings, Jörg-Christian Heinrich, Jean Herman, Ben Sprangers, Thierry Louat, Yves Moreau, Michael Schroeder

**Affiliations:** Biotechnology Center (BIOTEC), Technische Universität Dresden, 01307 Dresden, Germany; ESAT-STADIUS, KU Leuven, B-3001 Heverlee, Belgium; Interface Valorisation Platform (IVAP), KU Leuven, 3000 Leuven, Belgium; Laboratory of Molecular Immunology (Rega institute), KU Leuven, Belgium; Department of Pediatric Nephrology and Solid Organ Transplantation, University Hospitals Leuven, 3000 Leuven, Belgium; Department of Nephrology, University Hospitals Leuven, 3000 Leuven, Belgium; PharmAI GmbH, 01307 Dresden, Germany

## Abstract

Many drugs are promiscuous and bind to multiple targets. On the one hand, these targets may be linked to unwanted side effects, but on the other, they may achieve a combined desired effect (polypharmacology) or represent multiple diseases (drug repositioning). With the growth of 3D structures of drug-target complexes, it is today possible to study drug promiscuity at the structural level and to screen vast amounts of drug-target interactions to predict side effects, polypharmacological potential, and repositioning opportunities. Here, we pursue such an approach to identify drugs inactivating B-cells, whose dysregulation can function as a driver of autoimmune diseases. Screening over 500 kinases, we identified 22, whose knock out impeded the activation of B-cells. Among these 22 is the gene KDR, whose gene product VEGFR2 is a prominent cancer target with anti-VEGFR2 drugs on the market for over a decade. The main result of this paper is that structure-based drug repositioning for the identified kinase targets identified the cancer drug ibrutinib as micromolar VEGFR2 inhibitor with a very high therapeutic index in B-cell inactivation. These findings prove that ibrutinib is not only acting on the Bruton’s tyrosine kinase BTK, against which it was designed. Instead, it may be a polypharmacological drug, which additionally targets angiogenesis via inhibition of VEGFR2. Therefore ibrutinib carries potential to treat other VEGFR2 associated disease. Structure-based drug repositioning explains ibrutinib’s anti VEGFR2 action through a specific pattern of interactions shared between BTK and VEGFR2. Overall, structure-based drug repositioning was able to predict these findings at a fraction of the time and cost of a conventional screen.

## Introduction

The number of ways a drug can interact with its target is limited. Alex et al. estimate that there are 10000 interfaces [2] and Gao et al. reduce this number to 1000 distinct interface types [10]. While the exact number may be debatable, a limit itself has important consequences: there is redundancy and re-use in drug-target interaction, which makes it difficult to design a target-specific or one target-only drug. Hence, drug promiscuity, a drug hitting multiple targets, is rather the normality than the exception [16, 19, 27].

While drug promiscuity can be a source of a drug’s side effects, it can also be advantageous: In polypharmacology, drugs are designed to hit multiple targets associated to multiple desired effects in a disease. This ensures that the drug is more robust [3] to changes in the targets such as expression levels or mutation. Polypharmacology is successfully applied in practice. As an example, consider the kidney cancer drugs sunitinib, sorafenib, and pazopanib [23, 28], which are approved since 2006 and 2010, respectively. These drugs simultaneously act on proliferation and angiogenesis by binding (among others) both the platelet-derived growth factor receptors PDGFR and the vascular endothelial growth factor receptor VEGFR2. Targets such as PDGFR and VEGFR2 are particularly suitable to polypharmacology as they are kinases, the largest family of targets that bind to a common substrate adenosine triphosphate (ATP) [23].

Overington et al. argue that by nature of their close evolutionary relation and similar function, several members of the same family may be affected by a drug, which is thus promiscuous. In [19, 29], the authors quantify such polypharmacology by constructing a network of targets, which are related by shared drugs. The network is particularly dense for kinases, underlying the potential for side effects and polypharmacology. It also covers other classes of drug targets, such as e.g. G protein-coupled receptors (GPCRs), for which structural studies of selectivity and promiscuity [30, 34] pave the way for novel avenues in drug design through polypharmacology.

Besides polypharmacology, drug promiscuity can also support drug repositioning. Drug repositioning aims to identify novel indication for an existing drug. Often this is achieved by exploring diseases closely related to the original indication or by linking the known main drug target to a new disease. That latter was the case for Viagra with its repositioning from heart disease to erectyle disfunction. For both, old and new indication, the main target is the phosphodiesterase PDE5. However, repositioning is also possible if a drug hits two targets, one linked to the existing indication and the other to a new one. For example, the anti-herpes drug BVDU binds a viral thymidine kinase, but also the heat shock protein Hsp27, an anti-cancer target. Thus, BVDU’s promiscuity achieves two very different effects supporting its repositioning efforts [18] from herpes to cancer.

If drug promiscuity can be beneficial in polypharmacology and in drug repositioning, it becomes important to understand its source. As argued above, one reason for a drug to bind multiple targets is the limited space of binding interfaces [2, 16]. As a consequence, there are binding sites and interfaces which are very similar across unrelated targets and drugs. These similarities can be studied and uncovered algorithmically. In [24] the authors review target- and drug-centric approaches to polypharmacology. The former comprise alignments of binding sites and the latter fingerprints [7, 8], which characterize a drug and its interaction to a target. They can comprise chemical interactions such as hydrogen bonds, pi-stacking, or hydrophobic interactions together with geometric information on distances and angles. These interactions are encoded as standardized binary vectors, which become comparable. As a consequence, it is possible to search large repositories of drug-target complexes. Here, we will present and pursue such a structural approach to polypharmacology and drug repositioning with the goal of identifying candidate drugs for B-cell inactivation (see Fig 1).

**Fig 1.**
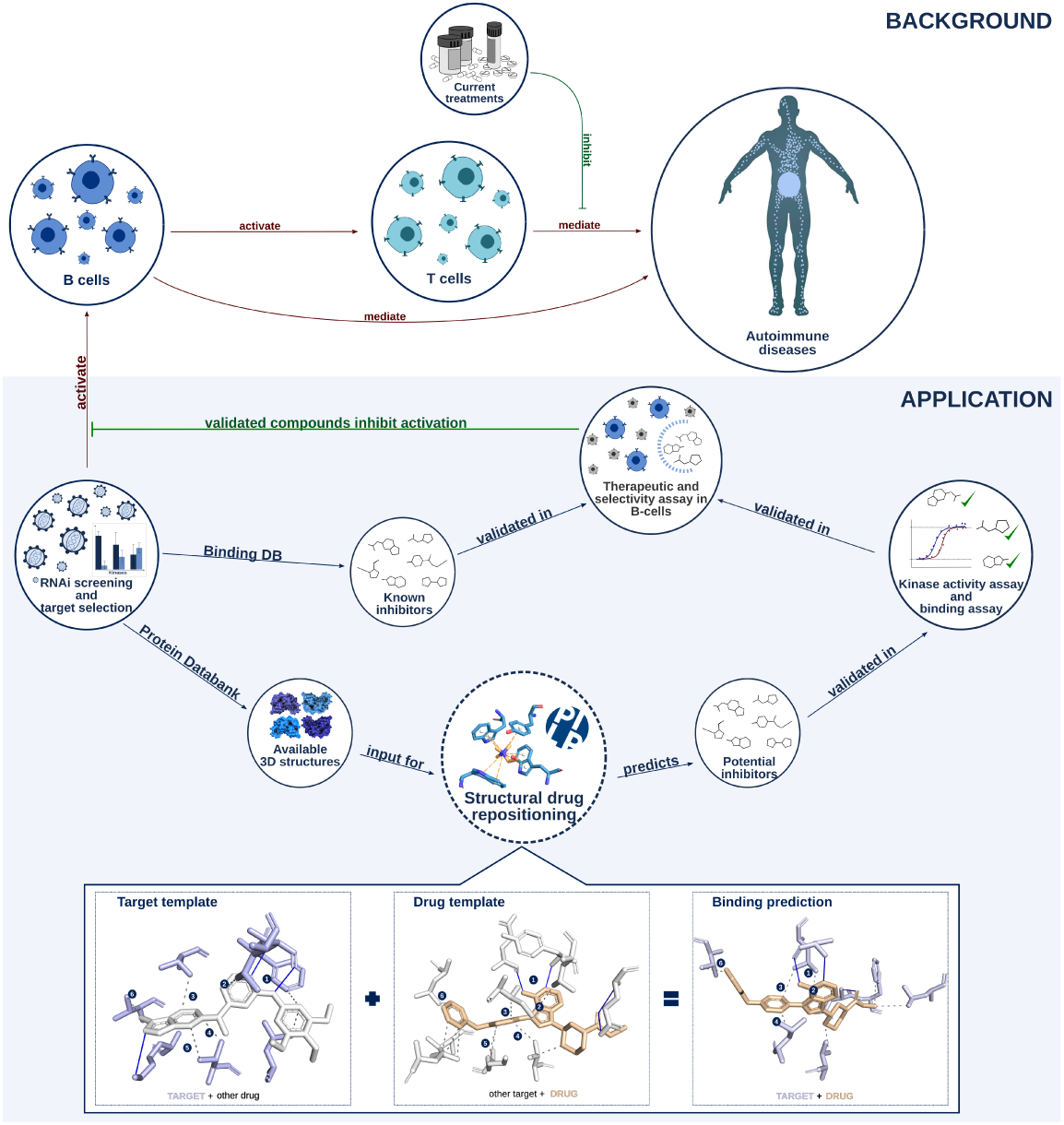
Overview of the study. Background. B- and T-cells play an important role in autoimmune diseases. Current treatments focus on inhibition of T-cell activation. Novel or improved treatments may emerge by inhibiting B-cells. **Application.** Through an RNAi screen, we identified kinases, that play a role in B-cell activation and are therefore suitable autoimmune targets. There are two routes to validate inhibitors against these kinases. First, known inhibitors may exist and second, structural drug repositioning may yield novel inhibitors after *in vitro* validation. Finally, all validated compounds are checked *in vitro* for selectivity and efficacy of B-cell inactivation.

B-cells produce antibodies and establish effective inter cellular communication for a controlled immune response through expression of cell surface markers and secretion of signaling molecules. B-cells play an important role in many blood cancers and in autoimmune diseases such as systemic lupus erythematosus, rheumatoid arthritis and psoriasis. Interestingly, there is a connection between cancer and psoriasis as angiogenesis, one of the cancer hall marks [14], and high expression of vascular endothelial growth factors (VEGFs) are present in psoriasis [13]. Subsequently, the polypharmacological cancer drugs pazopanib, sunitinib, and sorafenib lead to improvements in psoriasis [39]. This is particularly remarkable, as their initial indication is kidney cancer and not a blood cancer suggesting that blood cancer may be a repositioning opportunity. In fact, there are links between blood cancer and autoimmunity: Ibrutinib is used to treat leukemias and lymphomas as well as chronic graft vs. host disease. It was designed against Bruton’s tyrosine kinase (BTK), which are highly expressed in B-cells. Despite the designed specific inhibition of BTKs, there is recent evidence that ibrutinib is anti-angiogenic [31] suggesting that there are other targets.

Rituximab and bortezomib are two other cancer and autoimmune drugs, which act on B-cells. Rituximab is a monoclonal antibody, which binds to a surface marker of B-cells and then triggers cell-death [32]. Bortezomib is a cytotoxic small molecule widely used in cancer chemotherapy. Recently, it was tested for its autoimmune potential in lupus [1]. Both drugs kill B-cells rather than modulating their activity.

While there are links from cancer to auto-immune diseases for these drugs, they are nonetheless cytotoxic. Therefore, it is interesting to complement these drugs with novel small molecules, which inactivate B-cells and which have low toxicity. To this end, we will employ an assay to identify suitable targets driving B-cell inactivation and we will identify known and novel binders using a structural polypharmacology and drug repositioning approach. For target identification and lead validation, we will use a recently introduced assay [38], in which B-cell inactivation is read out via two cell surface markers. In [38], the authors compared six cell lines, thirteen stimuli, and eight read outs concluding that human Burkitt lymphoma cells (Namalwa) stimulated by the TLR9 ligand ODN2006 using expression-levels of the cell surface markers CD70 and CD80 is best suited to study B-cell inactivation. Cell surface markers such as CD70 and CD80 act as co-stimulatory molecules for the activation and differentiation of CD4+ T-cells. This is necessary for the development of an effective immune response. Furthermore, In vitro studies with murine B cells [37] and in vivo studies with murine disease models and transgenic mice [4] have indicated that markers CD70 and CD80 are also key molecules in the signalling for the regulation of B cell’s effector functions such as Ig production. Naive B-cells from patients with common variable immunodeficiency are markedly impaired in upregulating the costimulatory molecules CD80 and CD70 upon BCR cross-linking and the expression remained reduced even in presence of autologous helper CD4+ T cells. The insufficient upregulation of these two crucial costimulatory molecules could explain the poor class switching and, hence, reduced Ig serum levels, except for IgM [36].

## Materials and methods

The general methodology applied in this study is represented in Fig 1.Application and below there is a detailed description of all the step-by-step protocols performed in this work:

### RNAi screening and target selection

The approach developed, uses human Burkitt lymphoma Namalwa cells [26] stimulated with ODN2006. For RNAi screening, the MISSION LentiExpress TM shRNA library (Sigma-Aldrich, Diegem, Belgium) was used, which has 3 - 10 shRNA clones per kinase target. After transduction, cells were stimulated by ODN2006 for 24 hours. Cells were incubated with fluorescent antibodies for CD70 and CD80. After FACS sorting, expression was measured by amount of fluorescence. Then, the *y* -value was calculated as follows:

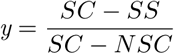

where *SC* is the expression of stimulated control, *SS* the stimulated sample, and *NSC* the non-stimulated control. Based on a close observation of the data, since generally, as observed, silencing of “hit” kinases gave a more potent inhibition of marker CD70 compared to marker CD80, clones with *y* > 0.8 for CD70 and *y* > 0.5 for CD80 were selected and resulted in the 22 kinase targets.

### Known inhbitors

Known inhibitors were retrieved from BindingDB in April 2018. A drug’s target in BindingDB had 100% sequence similarity to one of the 22 kinase targets and the Ki, Kd, or IC50 had to be less than 50*µ*M.

### Structural drug-repositioning

All ligand-target complexes in Protein Data Bank (PDB) [15] were represented as interaction fingerprints. Interaction fingerprints are binary vectors, where each feature represents a combination of interactions within an angle and distance range (see Fig 2). Interactions are obtained from PLIP [33] and comprise e.g. hydrogen bonds, hydrophobic contacts, pi stacking. Two interaction fingerprints designed by PharmAI company were compared by computing their Tanimoto score, i.e. the number of shared features divided by the number of overall features present in either vector. The score ranges from 0 (no features in common) to 1 (all features in common). For a set of fingerprints, the significance of a score is judged by p-value. When screening a template ligand-target complex against the PDB, the interaction fingerprint scores are sorted by p-value and only those with p-value less than 10^−3^ are retained. To apply structural drug-repositioning, we identified suitable templates in PDB. We found structures for 14 of the 22 kinase targets. This reduced to 10 after removing structures without interacting compounds or binding ions or biologically non-relevant small molecules according to BioLiP (zhanglab.ccmb.med.umich.edu/BioLiP). For these 10 kinases, there were a total of 141 drug-target templates.

**Fig 2.**
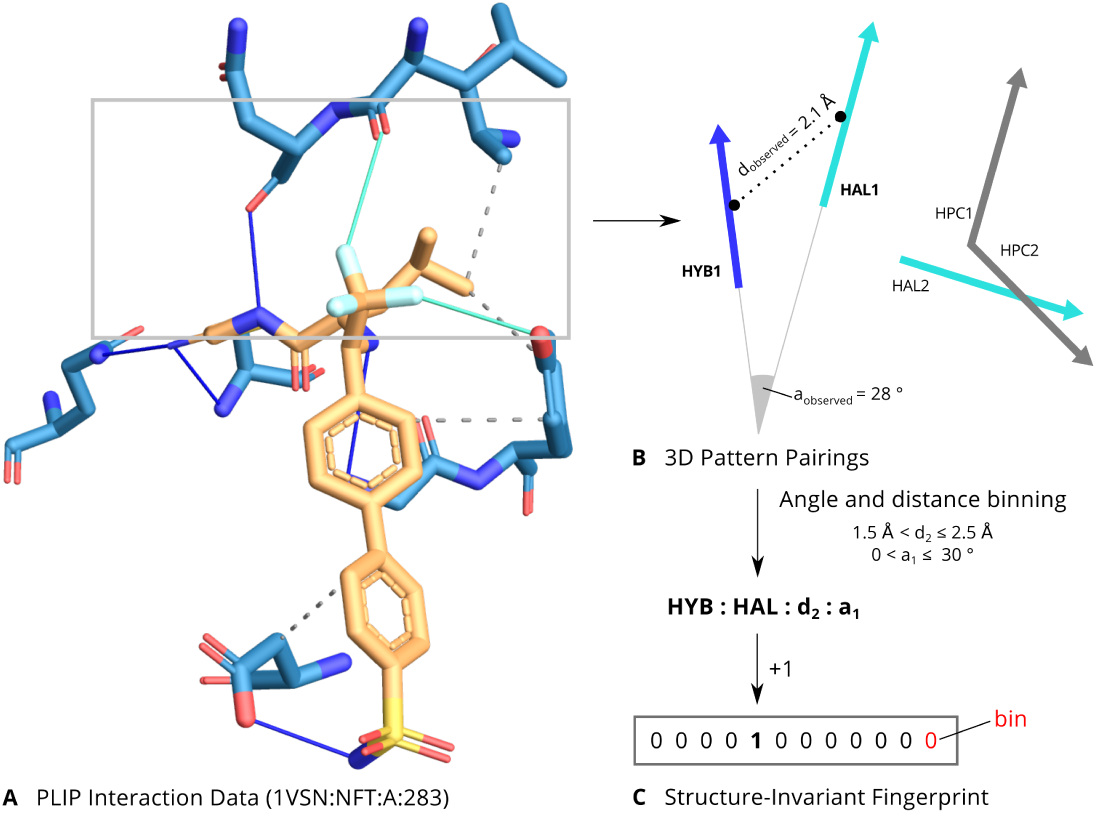
Structural drug-repositioning with PLIP interaction fingerprints. **A.**Schematic for the PLIP interaction fingerprints. Raw interaction data from PLIP are calculated for a complex. **B.**For each pair of interaction vectors, their distances and angles are calculated. **C.**Finally, the geometric measures are binned and combined with the types of the pairing into a string representation. Each such combination corresponds to a feature in the fingerprint, which is incremented for each observation of the encoded pairing. The resulting fingerprint is structure-invariant since it only considers the interaction vectors.

### Kinase activity test

Compounds predicted from structural drug repositioning were further filtered for FDA approval using DrugBank. Nutraceuticals, already known binders, and 6 compounds not available from Sigma-Aldrich were discarded. Dasatanib, a strong binder to all targets, was added as control. The 111 compounds were bought from Sigma-Aldrich and Selleckchem. Kinase inhibition was tested for them in duplicate at 1*µ*M and 10*µ*M concentrations resulting in 888 assays carried out by ReactionBiology. Data were normalized to percentage Enzyme Activity relative to DMSO controls. Compounds inhibiting more than 50 percent of the enzyme activity have been defined as strong inhibitors and inhibition between 30 and 50 percent as weak inhibition.

### Chemical structures and similarity

Chemical similarity was computed with the PubChem Score Matrix Service [22] with standard settings. Chemical structures were downloaded in SDF format. Heatmaps were plotted with the Heatplus package v2.16.0 in R 3.2.5 using hierarchical clustering on average distances.

### Links between targets and disease

To establish links between the 22 targets and disease we manually search PubMed (April-July 2018) and Open Targets [5] (December 2018). The search was done using the target name together with disease keywords cancer, cancer of the immune system, and autoimmune disease. We refined the latter with specific autoimmune disease such as lupus, multiple sclerosis, psoriasis, type 1 diabetes, Grave’s syndrome, inflammatory bowel disease, Hashimoto’s thyroiditis, Sjögren syndrome, celiac disease, rheumatoid arthritis. Cancer of the immune system was expanded with keywords myeloma, leukemia, lymphoma and cancer with tumour, adenoma, sarcoma, carcinoma, blastoma.

### Links between drugs and disease

PubChem was used (May 2018) to link drugs to diseases, which were grouped into the five high-level groups cancer, cancer of the immune system, autoimmune diseases, infection, and others.

### Drug-target-disease network

A drug-target-disease network was constructed by adding links from known as well as predicted and validated binders to their respective targets and adding the above target-disease and drug-disease links. The network was visualized with Cytoscape v3.6.0.

### Drugs’ known targets

The number of targets in Fig 7 was retrieved from BindingDB [12]. A target qualified if it is binding with a Ki, IC50 or Kd less than 50*µ*M. Targets are grouped by unique Uniprot Id.

### Kinase binding assay

Dose response curves for ibrutinib and suramin against VEGFR2 were determined with the KINOMEscan TM by DisvcoverX, a competition binding assay that quantitatively measures the ability of a compound to compete with an immobilized, active-site directed ligand. It is measured via quantitative PCR of the DNA tag linked to the kinase. An 11-point 3-fold serial dilution of each test compound was prepared in 100% DMSO at 100x of the final test concentration and subsequently diluted to 1x in the assay (final DMSO concentration = 1%). Most Kds were determined using a compound top concentration of 30000 nM. If the initial Kd determined was lower than 0.5 nM (the lowest concentration tested), the measurement was repeated with a serial dilution starting at a lower top concentration. A Kd value reported as 40000 nM indicates that the Kd was determined to be higher than 30000 nM. Binding constants (Kds) were calculated with a standard dose-response curve using the Hill equation:

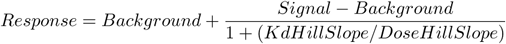

The Hill Slope was set to −1. Curves were fitted using a non-linear least square fit with the Levenberg-Marquardt algorithm.

### Interactions of Ibrutinib and suramin

Interactions in Fig 3.B were obtained from PLIP v1.4.0 [33] applied to PDB IDs 3cjg, 4ifg, 3c7q, 3gan.

**Fig 3.**
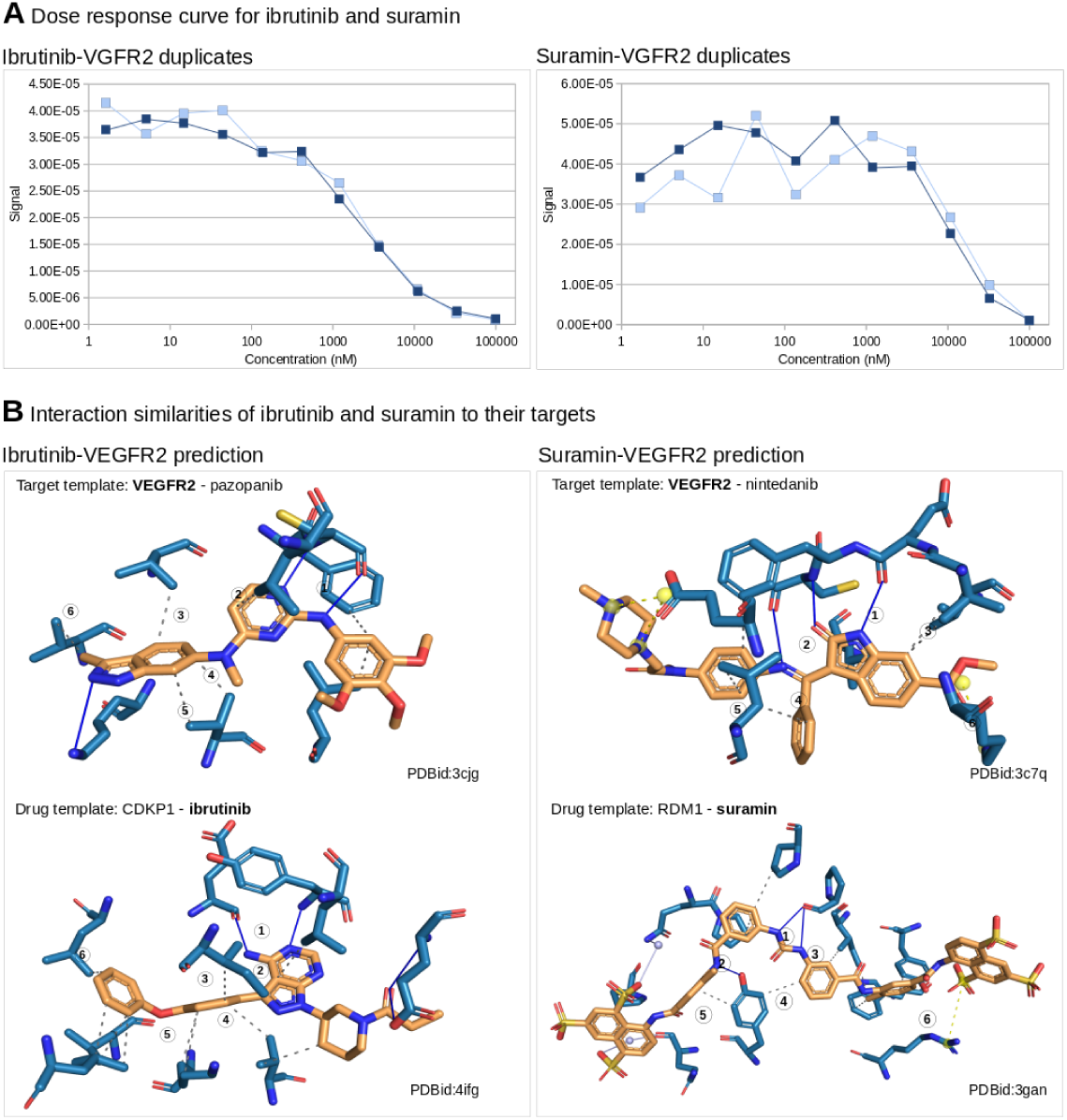
Ibrutinib and suramin are inhibitors of VEGFR2. **A.** Dose response curves for ibrutinib (KD value of 2*µ*M) and suramin (KD value of 25*µ*M) from the competition binding assay determined with the KINOMEscan TM by DisvcoverX prove that ibrutinib and suramin are *µ*M inhibitors of VEGFR2. **B.** Structural repositioning based in interaction of inhibitors (orange) to their targets (blue), predicts ibrutinib as VEGFR2 inhibitor since ibrutinib’s interaction to CDPK1 is similar to pazopanib’s interaction to VEGFR2. Both have (1) a double hydrogen bond (blue lines) and (2)(3)(4)(5)(6) hydrophobic interactions (gray dashed lines) in common. Similarly, suramin is predicted as VEGFR2 inhibitor. They have (1) a double hydrogen bond, (2) a simple hydrogen bond, (3)(4)(5) hydrophobic interactions and (6) a salt bridge (yellow dashed lines) in common. Subsequent in vitro validation proves both predictions correct.

### Therapeutic and selectivity index

In order to test the efficacy and selectivity of the hit compounds on B-cells, 4 assays were performed: B-cell assay, Mixed Lymphocyte Reaction (MLR), WST-1 assay on RPMI1788 cells (B-cell line) and cytotoxicity WST-1 assay on Jurkat cells (T-cell line). Compounds were prepared as 10 mM stock solutions in dimethylsulfoxide (DMSO)and were tested at 50-10-1-0.1-0.01 and 0.001 *µ*M in the 4 assays.

Blood samples of healthy volunteers were collected at the Red Cross of Mechelen, Belgium. Each donor consents to the use of his blood for research purposes. Human peripheral blood mononuclear cells (PBMCs) were obtained by density gradient centrifugation of the heparinized venous blood over Lymphoprep™ (Axis Shield PoC AS; density 1.077 ± 0.001 g/mL). The human B cell line RPMI1788 (American Type Culture Collection, USA) and the human T cell line Jurkat (European Collection of Cell Cultures, ECACC, England) were maintained in RPMI1640 culture medium (BioWhittaker®, Lonza, Verviers, Belgium) containing 10% foetal calf serum (FCS, HyClone® Thermo Scientific, United Kingdom) and 5 *µ*g/mL gentamicin sulphate (BioWhittaker®, Lonza, Verviers, Belgium) at 37°C and 5% CO2.

Highly purified naive peripheral human B cells were separated from fresh human PBMCs on magnetic columns by positive selection using CD19 magnetic beads according to the manufacturer’s instructions (MACS Miltenyi Biotech, Leiden, The Netherlands). The purity of the isolated naive B cells was ≥ 95% as analysed by flow cytometry. Freshly isolated human CD19+ B cells were then plated at 25 000 cells per well in a 384-well plate (Perkin Elmer, Zaventem, Belgium) in 55 *µ*L in DMEM medium containing 10% FCS and 5 *µ*g/mL gentamicin sulphate. After 7 days of stimulation with 0.1 *µ*M ODN2006 (InvivoGen, Toulouse, France), supernatant was taken for analysis of IgG using the AlphaLISA human IgG kit according to the manufacturer’s instructions (Perkin Elmer, Zaventem, Belgium). Analysis was performed with the EnVisionTM 2103 Multilabel Reader (Perkin Elmer, Zaventem, Belgium).

MLR assay (T cell assay): Freshly isolated human PBMCs and (responder cells) were resuspended in RPMI1640 medium containing 10% FCS and 5n*µ*g/mL gentamicin sulphate. RPMI1788 cells (stimulator cells) were inactivated by treatment with mitomycin C (Kyowa®, Takeda Belgium, Brussels, Belgium) for 20 min at 37 °C, washed and finally suspended in culture medium. An amount of 100 *µ*L of each cell suspension was mixed with 20 *µ*L of diluted compound. The mixed cells were cultured at 37 °C for 6 days in 5% CO2. DNA synthesis was assayed by the addition of 10 *µ*Ci (methyl-3H) thymidine (Perkin Elmer, Zaventem, Belgium) per well during the last 18 h in culture. Thereafter, the cells were harvested on glass filter paper and the counts per minute determined in a liquid scintillation counter (TopCount, Perkin Elmer, Zaventem, Belgium).

WST-1 assays on RPMI-1788 cells and Jurkat cells were performed as follows: 20 *µ*L of test compound dilution were added to each well containing exponential growing cells and the plates were incubated for 48 h at 37 °C, 5% CO2. Untreated cells and positive control (1% triton X-100, for the last 15 min) served as reference for maximum and minimum viability. At the end of incubation 100 *µ*L of supernatant were removed and replaced by 10 *µ*L of WST-1 solution (Cell Proliferation Reagent WST-1, Roche Applied Science). After 3 to 5 h incubation at 37°C, 5% CO2, optical density was measured at 450nm with the EnVisionTM 2103 Multilabel Reader.

B cell Therapeutic Index (B cell TI) was calculated as the ratio IC50 WST-1 assay RPMI1788/IC50 B cell assay; T cell Therapeutic Index (T cell TI) referred to the ratio IC50 WST-1 assay Jurkat/IC50 B CELL assay; while the selectivity index was calculated as the ratio IC50 MLR assay/IC50 B cell assay.

## Results

### Kinase targets for B-cell inactivation

As a first step towards novel B-cell modulators, we identified suitable kinase targets (see Fig 1.Application). To this end, human kinases were knocked out one by one and the knock out’s effect on cell survival and inactivation was measured using the assay introduced in [38]. Cell survival is important, as our goal is modulation of B-cells and not their depletion. As shown in Fig 4.A nearly all human kinases (501) were present in the RNAi library. After knock out, 400 proved to be non-lethal and 22 inactivate B-cells, i.e. the cell surface markers CD70 and CD80 were downregulated in expression (see Table 1).

**Table 1.** Targets identified in RNAi B-cell activation screen sorted by availability of structural data. The top ten targets are suitable for structure-based drug repositioning screening. For 10 targets (number 11 to 20) no structural were available and thus these target could not be used for structure -based drug repositioning. Silencing of all listed targets leads to strong the downregulation of CD70 and CD80 expressions in the B cell Namalwa cell line activated by the TLR9 agonist ODN2006 without causing cell death (column cell survival). A literature study links nearly all targets to cancer (C) and some to autoimmune diseases (AD) or cancer of the immune system (CIS). Overall, there are many novel autoimmune targets and KDR (VEGFR2) emerges as very promising target with 36 protein structures, potent decrease in expression of CD70 and CD80 following knock out of target in activated Namalwa cells without affecting cell viability and links to both cancer and auto-immunity.

|  | Gene name | Protein name | PDB structures | CD70 down regulation | CD80 down regulation | % Cell survival | C | CIS | AD |
| --- | --- | --- | --- | --- | --- | --- | --- | --- | --- |
| 1 | DAPK1 | Death-associated protein kinase 1 | 39 | 1.21 | 0.87 | 100 | x | x | x |
| 2 | **KDR** | **Vascular endothelial growth factor receptor 2 (VEGFR2)** | **36** | **1.27** | **0.98** | **75.6** | **x** | **x** | **x** |
| 3 | NEK2 | Serine/threonine-protein kinase Nek2 | 24 | 1.37 | 0.79 | 100 | x | x |  |
| 4 | CSNK1D | Casein kinase I isoform delta (CK1d) | 17 | 0.97 | 1.28 | 100 | x |  |  |
| 5 | EPHB4 | Ephrin type-B receptor 4 | 14 | 1.95 | 0.74 | 100 | x |  |  |
| 6 | CDC7 | Cell division cycle 7-related protein kinase | 4 | 1.88 | 0.83 | 100 | x |  |  |
| 7 | ILK | Integrin-linked protein kinase | 2 | 1.28 | 0.95 | 100 | x |  |  |
| 8 | MST-4 | Serine/threonine-protein kinase 26 | 2 | 2.63 | 0.73 | 100 | x |  | x |
| 9 | MAP2K2 | Dual specificity mitogen-activated protein kinase kinase 2 | 2 | 1.32 | 0.67 | 82.9 | x |  |  |
| 10 | MLK4 | Mitogen-activated protein kinase kinase kinase 21 | 1 | 1.81 | 0.54 | 99.7 | x |  |  |
| 11 | PXK | PX domain-containing protein kinase-like protein | 0 | 1.16 | 0.54 | 94.4 |  |  | x |
| 12 | PTK7 | Inactive tyrosine-protein kinase 7 | 0 | 1.07 | 0.71 | 86.2 | x | x |  |
| 13 | PAK3 | Serine/threonine-protein kinase PAK 3 | 0 | 2.83 | 0.98 | 86.8 | x |  |  |
| 14 | BRSK1 | Serine/threonine-protein kinase | 0 | 1.76 | 0.83 | 100 | x |  |  |
| 15 | FER | Tyrosine-protein kinase Fer | 0 | 1.60 | 0.95 | 100 | x |  |  |
| 16 | IRAK-3 | Interleukin-1 receptor-associated kinase 3 | 0 | 1.70 | 1.02 | 93.3 | x |  | x |
| 17 | MAP3K10 | Mitogen-activated protein kinase kinase kinase 10 | 0 | 1.65 | 0.93 | 87.0 | x |  |  |
| 18 | PRKAG2 | 5'-AMP-activated protein kinase subunit gamma-2 | 0 | 1.57 | 0.74 | 100 | x |  |  |
| 19 | SCYL1 | N-terminal kinase-like protein | 0 | 1.18 | 0.68 | 84.6 | x |  |  |
| 20 | STK33 | Serine/threonine-protein kinase 33 | 0 | 1.43 | 0.92 | 100 | x |  |  |
| 21 | TXK | Tyrosine-protein kinase TXK | 0 | 1.49 | 1.06 | 79.9 |  |  |  |
| 22 | WNK2 | Serine/threonine-protein kinase WNK2 | 0 | 1.47 | 0.86 | 100 | x |  |  |

The read out of CD70 and CD80 expression was crucial for this initial target identification. To quantify it, three steps must be considered: without stimulation, CD70 and CD80 expression in B-cells is low; upon stimulation, the markers are upregulated and finally, upon knock out of a kinase essential in B-cell activation, expression of the markers decreases. These two phases, the increase and the decrease, are compared to each other, i.e. it is not sufficient to compare expression levels of non-activated controls to activated knock outs. Instead activated controls have to be factored in. This is captured in the *y* -value (see Methods), which compares the markers’ upregulation after stimulation to their subsequent downregulation (or not) after knock out. Concretely, it is the fraction of expression level of activated control minus knock out and expression level of activated control minus non-activated control. A *y* -value close to zero indicates that the knock out does not influence B-cell’s activation, a value higher than one means that the knock out potentially inhibits activation of B-cells. Table 1 shows these fractions for the best clone of each of the 22 kinases. Besides the definition of the *y* -value, quality of knock out is important to consider. Per kinase to be silenced there are multiple RNA constructs, hitting different regions of the target gene, which may suffer from different off-targets. Due to this variability, there were between one and ten clones across the 22 kinases with varying degree of effectiveness in silencing. Subsequently, kinases were considered as suitable targets for B-cell inactivation/modulation if at least one clone shows *y* -value greater 0.5 for CD80 and greater 0.8 for CD70 (for more details check S1 File and S1 Fig)).

The 22 kinases, as shown in Table 1, were the starting point for the drug screen. As hinted in the introduction, drugs targeting B-cells are among others used in anti-cancer therapy and against autoimmune diseases. Therefore, we characterized the disease relation of the 22 identified targets focusing in particular on cancer, cancer of the immune system, and autoimmunity. A literature review revealed that nearly all targets are linked to cancer, four to cancer of the immune system, and five to autoimmunity (see Table 1). Among those five is VEGFR2, which is a key cancer target with approved drugs. Clinical trials are ongoing with exploring the inhibition of VEGFR2 as treatment for psoriasis (pazopanib, phase 1) and systemic scleroderma (nintedanib, phase 3). The other four targets with a link to autoimmunity are DAPK1 (inflammatory bowel disease), MST4 (Grave’s disease), PXK (systemic lupus erythematosus), and IRAK3 (rheumatoid arthritis) (for literature evidence check S1 Table).

To assess the 22 kinases’ potential as drug targets we consulted the Open Targets Platform [5], which covers over 20.000 targets with associated information on drugs and clinical trials. VEGFR2, EPHB4, CDC7, MAP2K2, and MAP3K10 are listed with VEGFR2 as most established target with a number of drugs already approved. Furthermore, we checked BindingDB [12], the largest database for drug-target binding affinities and found 22 drugs, each binding one or more of the 22 targets (see S3 Fig). 14 of the 22 kinases have at least one inhibitor in micromolar range and all but six drugs are polypharmacological drugs hitting multiple targets. In fact, 15 FDA approved drugs inhibit the same 13 kinases including VEGFR2, which documents the polypharmacological potential of these drugs. Most of these drugs are chemically closely related. Consider for example, erlontinib, vandetanib, and gefitinib (see Fig 5.A for their chemical structure), which belong to the same chemical class of quinazolines. To find a different compound with a better profile of toxicity and specificity we aimed to expand the scaffold space by running a structure-based drug repositioning screen.

### Structure-based screening to expand scaffold space

The starting point of a structure-based drug repositioning screen is a template drug-target complex, from which the chemical space beyond the drug’s scaffold and the given target’s fold is explored. A structure-based screening ignores the template’s global structure and instead focuses specifically on two aspects only: the binding site and the drug’s interaction to it (for more details see S2 File). These two features were extracted from the template and were compared to other drug-target complexes by aligning binding sites and scoring the similarity between the interactions (see Fig 2).

A prerequisite for structure-based screening is the availability of structural templates. Table 1 shows that 10 out of the 22 kinase targets have structural data available. For these 10 kinases targets there were a total of 141 templates. They were compared to over 300.000 ligand-target complexes in the Protein Databank PDB and ranked by p-value. The table also shows that there is huge variation from only one structure to over 30 for VEGFR2. Regarding candidate drugs, which may be predicted, PDB covers suitable structural data for 875 out of some 2500 FDA approved drugs. Based on these data, the structural drug repositioning screen predicts 157 approved drugs (see Fig 4.B). The 157 predictions are not equally distributed across the ten targets, but they correlate with the amount of structural data available for the target. e.g. for VEGFR2 with its 36 structures, there are 53 predictions, while MLK4 with one structure has three predictions only (for more details see S3 File).

**Fig 4.**
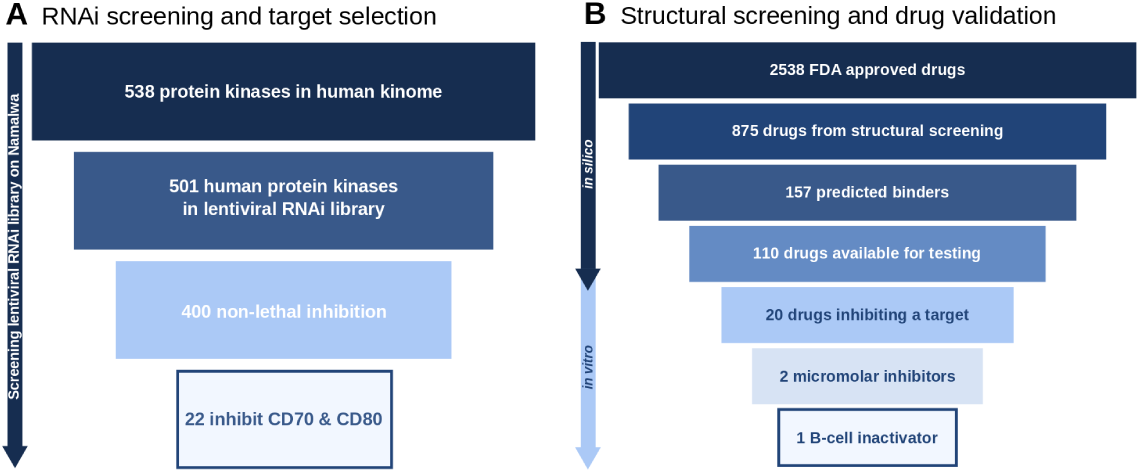
Selection of targets and inhibitors. **A.**From 538 human kinases to 22 involved in B-cell activation in Namalwa cells. Nearly all human kinases (501) are present in the RNAi library and for most of them (400) knock out is not lethal. Of those 400, 22 inhibit expression of both B-cell activation markers CD70 and CD80. **B.**From 2538 FDA approved drugs to 1*µ*M B-cell inactivator. Out of the 875 drug with available structures, the *in silico* screening yielded 157 positive binders. From the 110 drugs available for *in vitro* testing, 20 were validated as target inhibitors, 2 validated in kinase binding assay as positive micromolar inhibitors and finally one validated drug with positive effects in B-cells inactivation.

Subsequently these 110 drugs were tested at 10 *µ*M and 1 *µ*M on the panel of the 10 selected kinases (for more details see S4 File). We considered a 50% inhibition of the kinase activity as strong and 30 to 50% inhibition as weak. Table 2 summarizes the result. Only one compound, Ibrutinib, displays strong inhibition at 10 and 1 *µ*M on one of the kinase (VEGFR2); while 9 other compounds show strong inhibition at 10 *µ*M only. Ten additional compounds show weak inhibition at 10 *µ*M.

**Table 2.** List of new drug-target interactions predicted by interaction profile similarity and confirmed by experimental kinase activity assay. Enzyme activity was measured at a compound concentration of 1*µ*M and 10*µ*M. Drugs able to lower the enzyme activity to less than 50% (above dashed line) at 10*µ*M were defined as strong inhibitors while drugs that lowered kinase activity at 10*µ*M between 50% and 70% (below dashed line) were classified as weak inhibitors. Discarding the disinfectant Hexachlorophene because of its low medical interest, two compounds have been highlighted for showing inhibition at both 1*µ*M and 10*µ*M, ibrutinib and suramin.

| Compound | Target | % Enzyme activity<br>at 1μM | % Enzyme activity<br>at 10μM |
| --- | --- | --- | --- |
| Hexachlorophene | NEK2 | 84.92 | 00.02 |
| **Ibrutinib** | **VEGFR2** | **19.52** | **01.50** |
| Hexachlorophene | VEGFR2 | 109.5 | 05.14 |
| **Suramin** | **VEGFR2** | **72.67** | **06.48** |
| Toremifene | DAPK1 | 90.42 | 14.77 |
| Ibrutinib | DAPK1 | 95.59 | 23.72 |
| Saquinavir | DAPK1 | 87.09 | 27.81 |
| Tolcapone | CK1d | 91.69 | 35.58 |
| Ritonavir | DAPK1 | 92.58 | 38.46 |
| Etravirine | VEGFR2 | 108.4 | 41.08 |
| Thyroxine | NEK2 | 90.16 | 48.11 |
| Lopinavir | DAPK1 | 90.43 | 49.89 |
| Adenosine | CK1d | 87.72 | 50.22 |
| Diethylstilbestrol | NEK2 | 97.92 | 50.73 |
| Proflavine | NEK2 | 104.1 | 51.57 |
| Triiodothyronine | NEK2 | 111.9 | 55.67 |
| Adenosine | NEK2 | 96.13 | 58.99 |
| Triclosan | NEK2 | 99.17 | 64.62 |
| Triclosan | CDC7 | 70.45 | 65.43 |
| Propofol | EPHB4 | 66.96 | 68.44 |
| Prochlorperazine | NEK2 | 70.10 | 68.73 |
| Estradiol | DAPK1 | 110.6 | 68.98 |
| Amodiaquine | CK1d | 107.2 | 69.07 |
| Niacinamide | EPHB4 | 69.30 | 69.74 |

Thus, taking all analyses together, we have identified a total of 42 drugs (Fig 5), 22 known and 20 predicted and validated, which target at least one of the 20 target kinases and which thus carry the potential to inactivate B-cells. Before testing this potential, we summarize the chemical and disease space captured through these 42 drugs.

**Fig 5.**
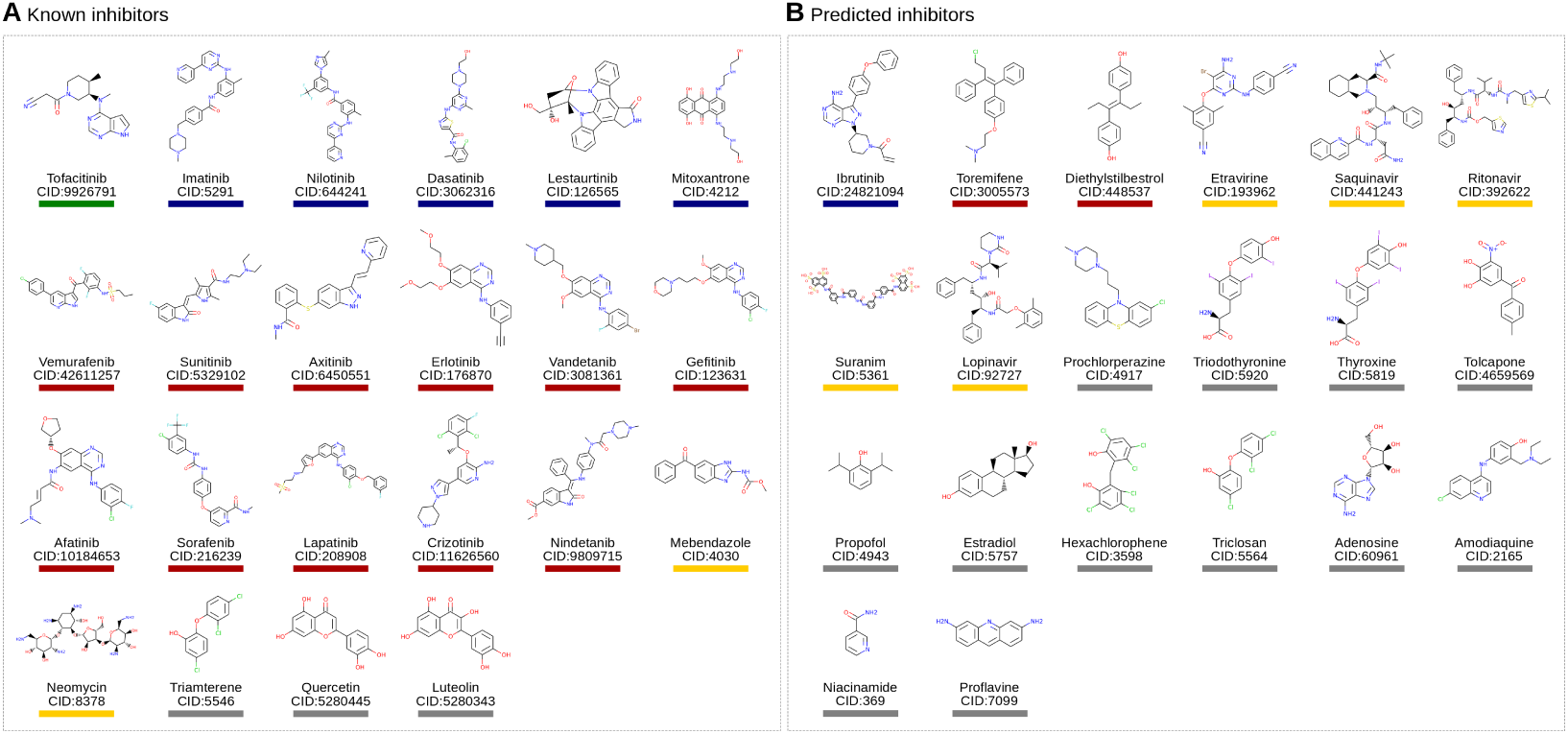
Chemical structures of known (A) and predicted (B) inhibitors. Drugs with drug name and compound ID (CID) are grouped by therapeutic indication: Autoimmune diseases(green), Cancer immune system (blue), Cancer (red), Infections (yellow), and others (gray).

### Chemical and disease space of identified drugs

Fig 6 shows a drug-target-disease network for the 42 drugs. The network depicts nicely that the majority of known drugs are used in cancer and are promiscuously targeting multiple targets. This is complemented by tofacitinib, an autoimmunity drug targeting many shared targets as well. In contrast, there is a sparse region in the network, where anti-infectives hit some of the targets including in particular VEGFR2. Ibrutinib is the exception to cancer drugs targeting only two targets.

**Fig 6.**
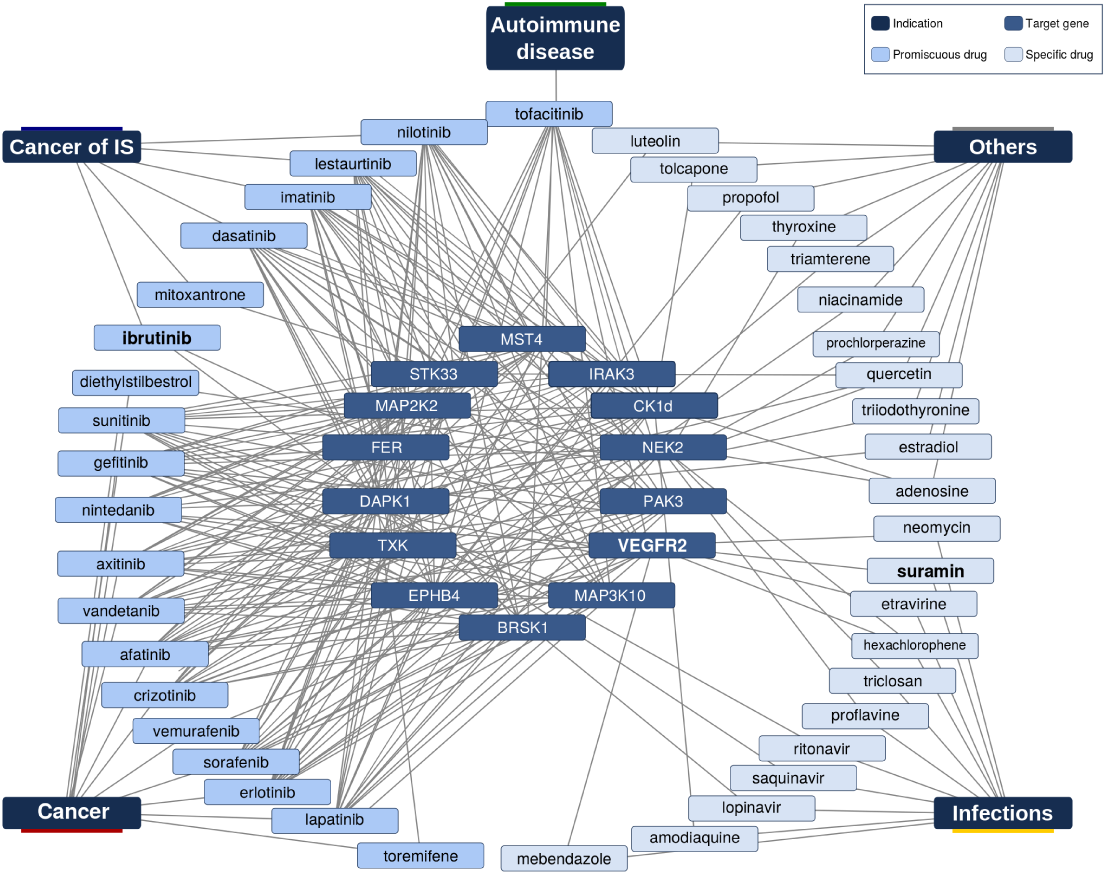
Drug-target-disease network. Inner circle (14 targets), mid circle (known and predicted inhibitors), outer circle (diseases). There is a tight network of cancer drugs and targets while there is only one known autoimmune compound, which binds, however, many of the cancer targets. VEGR2, the promising autoimmune target, we identified, is inhibited by cancer and anti-infection drugs.

The dense region of cancer drugs in the network may be due to their chemical similarity and resulting similar target profiles. Conversely, it can be expected that anti-infectives will have very different scaffolds. To understand this relation in detail, we compared the chemical structure of all 42 drugs against each other. Consider Fig 5 with the drugs’ structure. Visual inspection already reveals some similar scaffolds. As mentioned above, the anti-cancer drugs erlotinib, vandetanib, gefitinib, and afatinib are all quinazoline derivatives. To systematically assess these similarities we clustered the drugs by their chemical similarity according to PubChem’s CACTVS fingerprints [22]. CACTVS fingerprints represent chemicals as binary vectors of 880 dimensions. Each dimension represents a specific chemical feature. The chemical similarity of two drugs can then be defined as number of shared features divided by the overall number of features present in the two drugs, i.e. the Tanimoto score of the two fingerprints. The similarity can range between 0 (dissimilar) and 1 (similar).

Fig 7 depicts these chemical similarities for all pairs of the 42 drugs. Dark blue indicates high similarity. The dark blue group of four compounds in the top left quadrant represents the highly similar erlotinib, vandetanib, gefitinib, and afatinib mentioned above. This high-level view of the chemical compound space breaks down into the top left quadrant of similar compounds and the bottom right quadrant, which is completely dissimilar from the top left quadrant. To understand this grouping, we enriched the figure with additional information. On the bottom horizontal axis we labelled known binders in orange and novel binders in green. Broadly, the top left quadrant contains the known compounds, whereas the novel ones are the bottom right quadrant. This supports that the structural screen identified novel scaffolds not present in the known binders. Furthermore, we added to the vertical axis on the left the disease information for the drugs and on the right the number of targets. These bars show that the known binders in the top left quadrant have many targets and are anti-cancer, whereas the novel compounds in the bottom right cover various diseases and have few targets. At the border between known and novel quadrant is ibrutinib, an anti-cancer drug structurally close to many of the known drugs but not a known binder to any of the targets according to BindingDB. Furthermore, suramin, an anti-infective shows some weak structural similarity to the cancer drug mitoxantrone. The former has only very few targets, while the latter is highly promiscuous.

**Fig 7.**
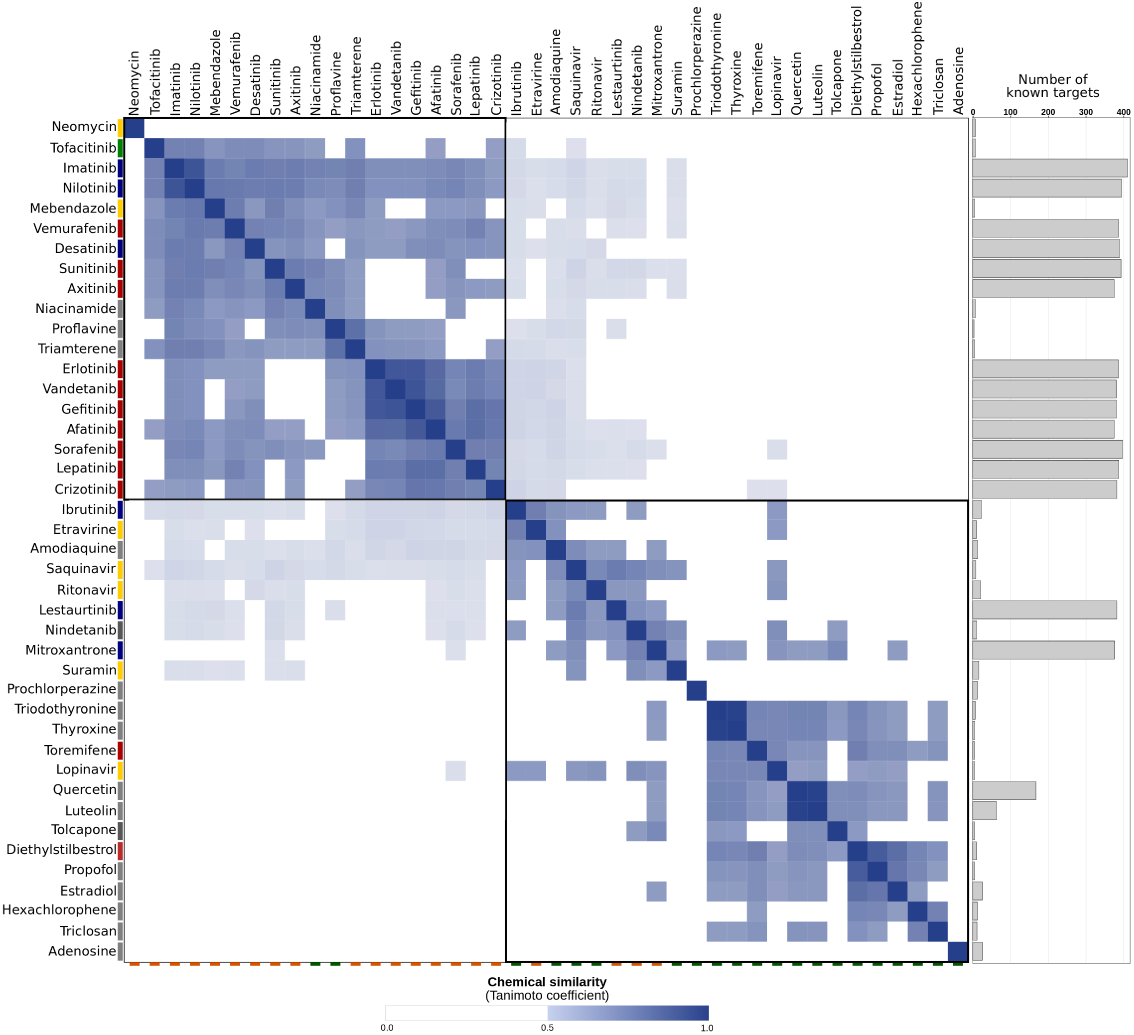
Pairwise chemical similarity of known and predicted inhibitors. Known inhibitors labeled with orange tag on bottom line and predicted ones with a green tag on bottom line. Dark blue boxes represent high chemical similarity of inhibitors (see Methods). Bars on the left are therapeutic indication (red=cancer, blue=cancer of the immune system, green=autoimmune disease, yellow=infection, and gray=other) and gray bars on the right represent the number of known targets (size of bar). The top left quadrant are mostly known inhibitors, which are chemically similar to each other and are mostly linked to cancer. The bottom right quadrant are new, predicted inhibitors, which have different scaffolds and are hence not derived from the known inhibitors. Consequently, these inhibitors have other therapeutic indications.

### Ibrutinib and suramin are micromolar VEGFR2 inhibitors

Ibrutinib and suramin emerged as promising leads from the kinase activity assay. The activity of ibrutinib on VEGFR2 is also confirmed by other experimental results in the LINCS kinomescan database [21]. To further sustain these findings, we carried out a dose response study. The curve in Fig 3.A plots drug concentration in nM against amount of bound kinase. Both experiments were carried out in duplicates. Ibrutinib binds to VEGFR2 at a Kd of 2*µ*M and suramin at a Kd of 25*µ*M.

To better understand how ibrutinib and suramin achieve this high affinity against VEGFR2, we checked their interaction profiles. The prediction of ibrutinib was based among others on ibrutinib’s interaction with CDPK1 (PDB 4ifg), which is very similar to pazopanib’s interaction with VEGFR2 (PDB 3cjg). This is a remarkable similarity, as CDPK1 is from the parasite Toxoplasma gondii and VEGFR2 from human. Despite two targets from completely different species, Fig 3.B shows that both interactions comprise a double hydrogen bond (1) and five hydrophobic interactions in similar position (2), (3), (4), (5), and (6). In a similar way, suramin’s binding to RDM1 (PDB 3gan) is similar to nintenadib’s interaction with VEGFR2 (PDB 3c7q). Again, this similarity is very remarkable, as RDM1 is a regulator of DNA methylation in Arabidopsis thaliana and VEGFR2 is from human. Nonetheless, both comprise in their ligand interaction a double hydrogen bond (1), a simple hydrogen bond (2), three hydrophobic interactions (3), (4), (5) and a salt bridge (6).

These examples not only document scaffold hopping from pazopanib to ibrutinib and from nintenadib to suramin, but also target hopping from parasitic CDPK1 to human VEGFR2 and from plant RDM1 to human VEGFR2. These jumps in target space are large. They cross family and species boundaries. As such they illustrate the limited space of binding sites, which was eluded to in the introduction.

### Ibrutinib inactivates B-cells

In a final validation step, we tested the drugs’ ability to impede B-cell’s activation (for more details see S5 File). We wanted inhibition of activation without cell death and specific to B-cells. We introduce the therapeutic index, which relates toxicity to inactivation. We calculate it for B- and T-cells. From this we compute the specificity index as ratio of B-cell to T-cell therapeutic index. Our goal is to uncover compounds with high B-cell therapeutic index and high specificity index. Indices are computed from duplicates. For details on cell lines used see Methods section.

As a main result, ibrutinib has a very high B-cell therapeutic index of over 40000 confirming ibrutinib’s ability to inactivate B-cells at low toxicity. A selectivity index of over 5000 also means that ibrutinib interferes only little with T-cells. Table 3 summarize these indices including 15 other compounds. For suramin, the therapeutic and specificity indices are close to one and hence suramin is not a B-cell modulator, although it is a micromolar VEGFR2 inhibitor. Tofacitinib, which can be seen as a control, due to its approval for autoimmune indications, has a good B-cell therapeutic index but lacks specificity.

**Table 3.** Potency on B-cell, B-cell therapeutic index (TI), T-cell therapeutic index, and specificity index (SI). The B cell IC50 is given in *µ*M and is the mean of 4 independent experiments. The therapeutic index relates toxicity to activity. B cell Therapeutic index (B cell TI) is the ratio between the toxic concentration and active concentration on B-cell and was calculated as IC50 WST-1 assay RPMI1788/IC50 B-cell assay. T cell Therapeutic index (T cell TI) is the ratio between toxic and active concentrations on T-cells and was calculated as IC50 WST-1 assay Jurkat/IC50 B-cell assay. The selectivity index is the ration between active concentrations on T-cells and B-cells and was calculated as IC50 MLR assay/IC50 B-cell assay. A high value of Therapeutic Index corresponds to high activity at low toxicity and is hence desirable. A high value of Selectivity Index indicates high specificity to B-cells rather than T-cells. Ibrutinib has high B-cell TI and SI.

| Drug | B-cell<br>IC50 ($\mu\text{M}$) | B-cell<br>TI | T-cell<br>TI | T/B-cell<br>SI |
| --- | --- | --- | --- | --- |
| **Ibrutinib** | **0.0002** | **3856** | **81090** | **5530** |
| Dasatinib | 0.046 | 0,71 | 160,0 | 0,07 |
| **Tofacitinib** | **0.432** | **>115,6** | **>115,6** | **0,61** |
| Proflavine | 0.59 | 5,14 | 8,55 | 7,50 |
| Amodiaquin | 0.66 | 42,33 | 39,43 | 14,73 |
| Hexachlorophene | 5.34 | 0,79 | 1,10 | 0,27 |
| Etravirine | 5.90 | 1,35 | 5,15 | 1,23 |
| Prochlorperazine | 6.12 | 1,67 | 3,52 | 0,76 |
| Triclosan | 20.88 | 1.32 | 1,25 | 1.64 |
| Diethylstilbestrol | 31.07 | 1,01 | 0,22 | 0,59 |
| Estradiol | 33.91 | 1,06 | 0,58 | 0,62 |
| Thyroxine | 34.10 | >1,47 | >1,47 | >1,47 |
| **Suramin** | **35.20** | **>1,42** | **>1,42** | **0,84** |
| Thyronine | 36.82 | >1,36 | >1,36 | 1,20 |
| Adenosine | >50 | 1,00 | 1,00 | 1,00 |
| Nicotinamide | >50 | 1,00 | 1,00 | 1,00 |

## Discussion

Drug promiscuity is a common phenomenon and we have exploited it here in a structure-based drug repositioning screen to identify ibrutinib, a novel VEGFR2 inhibitor and a B-cell inactivator.

VEGFR2 is a receptor-type tyrosine kinase (RTK) with a key role in the regulation of angiogenesis and lymphangiogenesis, which makes it an attractive target for antiangiogenic and proangiogenic therapies [35]. A lot of VEGFR2 neutralizing agents have been developed for cancer treatment, such as the human antibody bevacizumab (Avastin) [35], and the kinase inhibitors sunitinib, sorafenib, axitinib and vandetanib. This kinase showed already an interesting connection with the autoimmune disease psoriasis [17], and two of its inhibitors have been repositioned from the anti cancer indication to immunomodulation and are currently in clinical trials for the treatment of psoriasis (pazopanib) and systemic scleroderma (Nintedanib).

While this discovery of a confirmed binder of a new scaffold proves the power of the in silico approach, it raises the question of the approach’s general applicability. In drug repositioning, one can take a target- or a drug-centric approach. In a target-centric approach, the target of the repositioned drug is the key. It was associated with one disease and is now found to be relevant in another disease. Hence, the drug may be repositioned from the first to the second disease. In a drug-centric approach to drug repositioning, the prediction of a novel target, which is linked to a disease, forms the base for repositioning. In this work, we pursued both approaches.

The RNAi screen on the human B-cell line Namalwa identified novel B-cell targets by using CD70 and CD80 expression as read-out for B-cell activity. We must clarify that the role of CD70 and CD80 is not exclusive of B-cells activity, and therefore their inhibition might also be important in contrasting T-lymphocytes over-activation with an advantageous pleiotropic effect in immunosuppressive therapy. Among the various stimuli investigated for B-cell activation, in vitro stimulation with TLR9 agonist ODN2006 resulted in the broadest phenotypic change profile in human polyclonal B-cells. Compared with an assay of human B-cell IgG, which checks the level of IgG production by primary B-cells after 7 days incubation with the same stimulation agent ODN2006, the Namalwa-ODN2006 protocol takes only 24 hours, making it ideal for a fast first line screening. Furthermore, the heterogeneity between the different human blood donors, the heterogeneity after isolation and purification from peripheral blood, the limited yield of the purification and the short longevity make primary B-cells less optimal for repeatable assays. So a monoclonal population of dividing cells such as the human B-cell line Namalwa stabilizes the assay by overcoming the aforementioned limitations.

Our analysis of known binders to these targets is target-based drug repositioning. It was fruitful as many of the targets were established drug targets with associated approved drugs. In fact, our analysis showed that these drugs are highly promiscuous and hit many of the targets, which is possibly an intended polypharmacological design. Their chemical analysis showed highly similar compounds of the same scaffolds. Therefore, we applied structure-based screening to implement scaffold hopping away from these known binders into novel, uncharted chemical space. This second approach was drug-centric, as the structure-based screen across the Protein Databank PDB led to novel drug-target predictions. The most prominent ones were suramin and ibrutinib, both binding VEGFR2. If we consider this repositioning at a higher level, then suramin may be moved from infectious disease (sleeping sickness and river blindness) to cancer and ibrutinib from cancer to autoimmunity. While both drugs suffer from side effects, they are nonetheless serious drug repositioning candidates: suramin is tested in a phase I/II study against autism (see clinicaltrials.gov) and the FDA approved ibrutinib in 2017 for host-vs-graft disease.

While the above results focus on the most established target, VEGFR2, the RNAi screen generated another 21 novel B-cell targets overall. For some there is a link to autoimmunity and for a few there are drugs known to inhibit them. However, future work should fully explore all candidates regarding their relation to disease. This is difficult for any target and any drug: Even if a drug and a target can be linked to a disease, it still does not necessarily mean that the drug acts through the target. As an example, the RNAi screen suggests that silencing of VEGFR2 leads to B-cell inactivation, the inhibition of VEGFR2 with ibrutinib confirmed this effect, whereas inhibition with suramin did not. Generally, besides a methodological difference in silencing and inhibition, ibrutinib may have a more favorable polypharmacological profile for B-cell inactivation than suramin has. In a nutshell, both target- and drug-centric drug repositioning can be successful. The latter has the advantage of moving to novel scaffolds. And last, but not least, the structure-based drug repositioning does not only comprise the prediction, but also a transparent physio-chemical explanation for the prediction. Thus, giving confidence and a starting point for subsequent optimization steps.

A prerequisite for these successes is the availability of structures available in the Protein Data Bank (PDB). While there are far less structures than sequences in public databases, the growth of structural data is none the less impressive: PDB has more than doubled in size over the last seven years. Today, it contains 3D structures of over 1200 different drug targets and more than 60% of all PDB structures contain proteins in complexe with biologically relevant ligands. For the present screen, PDB contained suitable structures for about half of the targets. S2 Fig shows how these structures became available over time. It reveals that 15 years ago, structure-based drug repositioning would not have been possible for the identified 22 targets. Assuming a continuing linear growth, the figure suggests that in another 15 years the complete set of targets can be surveyed. Thus, the potential of structure-based drug repositioning is visible today and will further increase over time. Technological advance is also likely to speed up progress, since improvements in structure prediction and modelling may soon reach the maturity to close the structural gap faster.

The structural drug repositioning screen as carried out here considered two aspects: the ligand-target interaction and superposition of binding site and ligand. Generally, many approaches in the past have focused either on the ligand (see e.g. [6]) or on the target (geometrical and/or chemical similarities of the target proteins [20]). Our approach combined both views. The quality of such an integrated approach can be measured by the hit rates achieved. Here, we considered 110 compounds after prediction and filtering steps. For 15 of these prediction there was experimental evidence of high affinity binding in BindingDB and for other 20 unknown binders our kinase inhibition assay provided confirmation. This corresponds to a very high hit rate of 32%.

Such hit rates are unprecedented in traditional high-throughput screening, but they come at a cost. In classical high-throughput screening, there is a large compound library, which may be already focused on a specific type of compounds. In contrast, the structural screen is limited by the compounds in PDB. Overall, there are over 30000, which is substantial. But in comparison to the million of compounds available in databases such as PubChem it is small. This gap in size is reduced if one adds additional selection criteria. In drug repositioning, this can be e.g. the approval of the compounds. There are some 2500 approved drugs and for the structural screening some 800 are available in PDB. So, the orders of magnitude in size difference reduce to a factor of three. A possible disadvantage of a smaller library may be the inability to cover a large chemical space. However, this problem does not arise in the structural screen as carried out here, since the focus on drug-target interaction supported scaffold hopping. It ensured that huge steps in chemical space were possible and were indeed observed in the resulting data. Finally, *in silico* screening implies low cost and time. Instead of 8750 validation experiments (875 approved drugs with structure times 10 targets), there were only some 100, a cost reduction by nearly two orders of magnitude. In summary, the combination of a small in silico library with the algorithmic approach ultimately leads to a very high hit rate over a large chemical space at a low screening cost.

Other computational approaches also pursue *in silico* drug-target prediction by using text mining, machine learning or chemical similarities, among other methods. When comparing our method to the three most cited approaches [9, 11, 25], in order to check whether they are able to predict the ibrutinib-VEGFR2 connection or not, we came out with the following: SwissTargetPrediction (129 citations) is based on similarity measures of chemical structure and molecular shape. Performing a target screening for Ibrutinib, it predicted a long list of TYR kinases, including EGFR, but no specific link to VEGFR2. On the other hand, SuperPred (98 citations) is based on structural fingerprint similarities and text mining for target prediction. It identifies no known targets for Ibrutinib and while it is able to predict 5 TYR kinases with good E values, non of them is a direct link to VEGFR2. Finally, Hitpick (45 citations) combines two methods (1-Nearest-Neighbour and machine learning) in order to calculate 2D similarity. For ibrutinib, it predicted only 2 TYR kinases as targets, but non of them VEGFR2.

Although they all successfully predicted targets from the Receptor protein tyrosine kinase class, which is the same of VEGFR2, none of them was able to actually make the direct link as our method did. Furthermore, all of the mentioned approaches are based in 2D/3D similarity measures of the compounds and in available assay data coming from databases. Therefore, their screening results often in similar scaffolds and protein classes, being unable to make such specific prediction for distant biochemical entities.

## Conclusion

B-cells play an important role in diseases ranging from auto-immunity to cancer. There is an unmet need for small molecules inactivating B-cells. Here, we present ibrutinib as such a candidate immuno-modulator. In a first step, we ran an RNAi screen with a recently introduced drug screening assay for B-cell inactivation. We found 22 kinases, whose silencing inactivates B-cells. Many of these targets are already drug targets with approved drugs, which will be beneficial for future drug repositioning efforts. We focused specifically on VEGFR2, as it is already an established cancer drug target and there is evidence for a role in autoimmunity, too. For lead identification, we pursued two approaches. We evaluated existing drugs affecting the identified targets. As we found mostly chemically similar cancer drugs with a polypharmacological design hitting many of our targets, we wanted to expand the scaffold space. We used structural drug repositioning to predict novel inhibitors, which we then validated in an enzyme activity and binding study. We identified the cancer drug ibrutinib and the anti-infective suramin as micromolar VEGFR2 inhibitors. In a final step, we tested B-cell inactivation and found ibrutinib as a very effective inhibitor with very high therapeutic index and good specificity to B-cells rather than T-cells. The discovery is particularly remarkable as the underlying scaffold hopping is tied to target hopping. At the source of the prediction are similarities in the binding sites of human VEGFR2 to a parasitic and to a plant protein. These similarities are remarkable as they are unlikely of an evolutionary origin since not only family boundaries are crossed, but also species boundaries. Instead, they document the large, but limited number of binding sites, as postulated in [2, 10]. Overall, the in silico structural screen produced results at a very high hit rate, which translates into low cost and time. It was able to predict chemically distant compounds thus covering a large chemical space.

## Supporting information

RNAi screening

Compounds screening

Protein-drug predictions

Activity assay

Bcell assay

RNAi screening for target identification

Available structures for 22 targets over time

## Supporting information

**S1 File. RNAi screening.** Results of the shRNA transfection of human Burkitt lymphoma cells (Namalwa) stimulated by the TLR9 ligand ODN2006. Per each clone the name of the silenced gene (‘Gene’ column), the number of Namlwa cells aquired by the FACS (‘Cell number’), the percentage of inhibition of over-expression of surface proteins CD70 (‘% inh CD70’) and CD80 (‘% inh CD70’) are reported.

**S2 File. Compounds screening.** Dataset of 157 compounds resulting from the computational screening based in interaction similarities. It contains the compound name, compound id (CID), inchikey, drug category if possible, binding evidence coming from Binding DB, related diseases coming from PubChem, best p-value and finally individual P-values for each compound-targets prediction.

**S3 File. Protein-drug predictions.** List of 1828 predictions between the 157 compounds and the 22 targets. It contains the compound id (CID), the protein target, the complex id (pdbid:hetid:chain:position) of the template, the complex id of the hit, the method making the prediction and the p-value of the prediction.

**S4 File. Activity assay.** Results of the kinase activity assay performed for the 111 purchased compounds. It contains the compound name, the target name and the enzyme activity value of 2 data points at 1 and 10 µM.

**S5 File. Bcell assay.** Results of the efficacy and cytotoxicity tests of 16 of our kinase inhibitors (‘Compound’ column) on B-cell and T-cell lines. The four type of assays are reported: ‘B-cell efficacy’, ‘T-cell efficacy’, ‘Cytotoxicity/ WST1 RPMI 1788’ and ‘Cytotoxicity/ WST-1 Jurkat’. For each assay the mean of 4 different independent measures and the standard deviation (SD) are reported. Three different indexes have been calculated: ‘Therapeutic Index (TI)/ WST1 RPMI 1788/B-cell’ (B-cell/B-cell), ‘Therapeutic Index (TI)/ WST1 Jurkat/B-cell’ (T-cell/B-cell), and ‘Selectivity Index (SI)/ MLR/B-cell’ (T-cell/B-cell). Mycophenolate Mofetyl and Cyclosporine A have been used as positive controls.

**S1 Fig. RNAi screening for target identification.** The level of inhibition of upregulation (y), of CD70 (dark blue dots) and CD80 (light blue dots) for each clone of the selected genes (x axe) have been plotted plotted. The clones laying in the bottom part of the graph with y ¡ 0 (red part), showed an expression of the surface receptor (Ssample or Ss on the side bar) higher than the stimulated control cells (Sctrl or Sc); the clones (Ss) with 0 ¿ y ¿ 1 (yellow part) had a level of CD70/CD80 expression lower than the stimulated controls (Sc) but higher than the non-stimulated control cells (NSctrl) or NSc); the clones (Ss) placed in the area with y ¿ 1 (green part) showed an exprssion of the activation markers lower also than the non-stimulated controls NSc. Those genes have been selected for showing the y of at least one clone above both the threshold of 0.8 for CD70 (dark blue dotted line) and 0.5 for CD80 (light blue dotted line).

**S2 Fig. Available structures for 22 targets over time.** Over the last 15 years, structures for half of the identified target kinases were deposited in PDB, so that today there is sufficient data for structure-based drug repositioning available. Before the year 2002, this type of screening would not have been possible. In the future, it will further improve.

**S3 Fig. Known drugs bind unspecifically.** Known drugs binding 22 identified kinases with Ki, Kd, or IC50 ¡50µM. Drugs are colored by indication: red for cancer, blue for cancer of immune system, green for immunomodulation, yellow for infection and gray for other indications. Overall, the majority of drugs are anti-cancer and bind unspecifically.

**S1 Table. Literature evidence for a target’s association to disease.** Links in litterature between each gene target (‘Target’ column) and some pathological conditions (‘Disease’ column) such as Cancer, Tumors of Immune System (Lymphoma, Leukemia and Multiple Myeloma), and Autoimmune Diseases (Inflammatory Bowel Disease, Psoriasis, Lupus Erythematosus, Graves’ Disease and Rheumatoid Arthritis) are reported (‘PMID’ column).

## Acknowledgments

Thanks to Florian Kaiser and Arnout Voet for valuable feedback.

## Author contributions statement

MA, DP, MS conceived the study, generated and analysed data, and wrote the manuscript. SS, VJH, MA implemented structural screening. GJ, JCH, JH, KvB, BS carried out experiments and analysed data.

