## Supplementary figures and images for "Structure-based drug repositioning explains ibrutinib as VEGFR2 inhibitor"

### Available structures for 22 targets over time

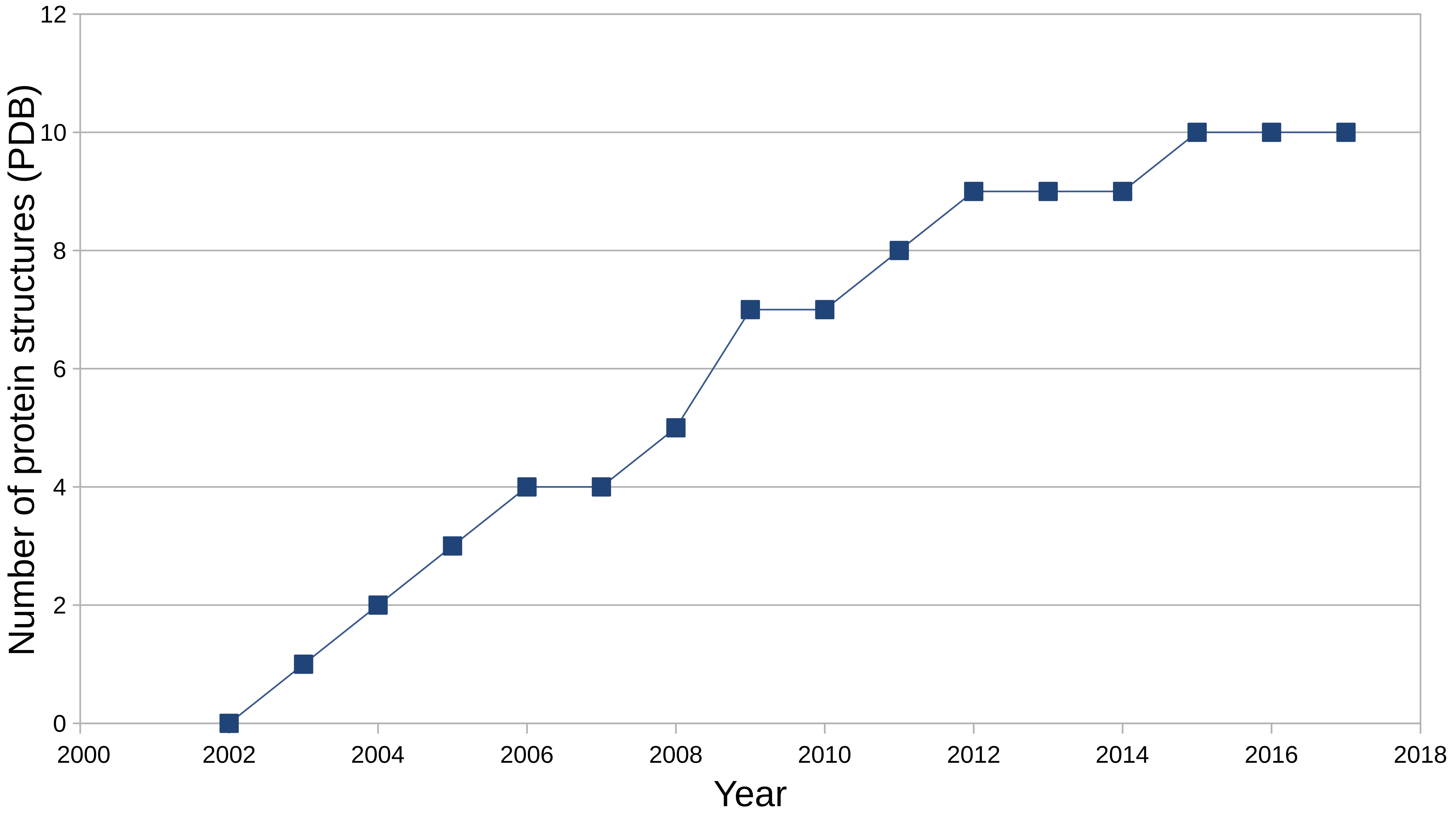

### RNAi screening for target identification

#### Kinase inhibition in CD70 and CD80

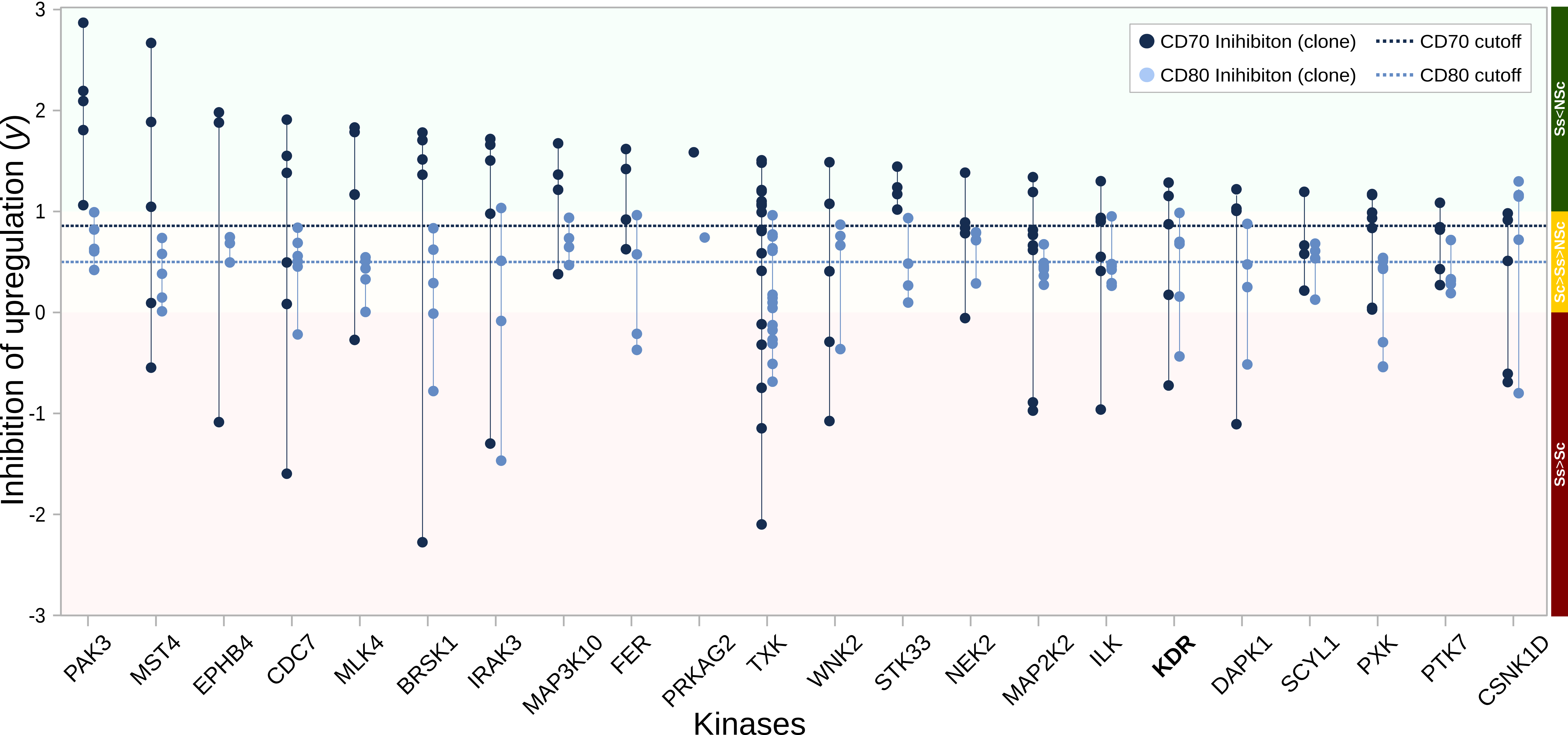
