## Supplemental Data 2 for "Structure-based drug repositioning explains ibrutinib as VEGFR2 inhibitor"

| Drug \ Target | VEGFR2 | DAPK1 | CK1d | NEK2 | EPHB4 | MAP2K2 | MST4 | STK33 | TXK | IRAK3 | BRSK1 | MAP3K10 | PAK3 | FER | WNK2 | PRKAG2 | SCYL1 | PXK | PTK7 | ILK | CDC7 | MLK4 |
|---|---|---|---|---|---|---|---|---|---|---|---|---|---|---|---|---|---|---|---|---|---|---|
| Imatinib | ✓ | ✓ | ✓ | ✓ | ✓ | ✓ | ✓ | ✓ | ✓ | ✓ | ✓ | ✓ | ✓ | ✓ | | | | | | | | |
| Dasatinib | ✓ | ✓ | ✓ | ✓ | ✓ | ✓ | ✓ | ✓ | ✓ | ✓ | ✓ | ✓ | ✓ | ✓ | | | | | | | | |
| Erlotinib | ✓ | ✓ | ✓ | ✓ | ✓ | ✓ | ✓ | ✓ | ✓ | ✓ | ✓ | ✓ | ✓ | ✓ | | | | | | | | |
| Gefitinib | ✓ | ✓ | ✓ | ✓ | ✓ | ✓ | ✓ | ✓ | ✓ | ✓ | ✓ | ✓ | ✓ | ✓ | | | | | | | | |
| Nilotinib | ✓ | ✓ | ✓ | ✓ | ✓ | ✓ | ✓ | ✓ | ✓ | ✓ | ✓ | ✓ | ✓ | ✓ | | | | | | | | |
| Sunitinib | ✓ | ✓ | ✓ | ✓ | ✓ | ✓ | ✓ | ✓ | ✓ | ✓ | ✓ | ✓ | ✓ | | | | | | | | | |
| Crizotinib | ✓ | ✓ | ✓ | ✓ | ✓ | ✓ | ✓ | ✓ | ✓ | ✓ | ✓ | ✓ | ✓ | | | | | | | | | |
| Vandetanib | ✓ | ✓ | ✓ | ✓ | ✓ | ✓ | ✓ | ✓ | ✓ | ✓ | ✓ | ✓ | ✓ | | | | | | | | | |
| Sorafenib | ✓ | ✓ | ✓ | ✓ | ✓ | ✓ | ✓ | ✓ | ✓ | ✓ | ✓ | ✓ | ✓ | | | | | | | | | |
| Tofacitinib | ✓ | ✓ | ✓ | ✓ | ✓ | ✓ | ✓ | ✓ | ✓ | ✓ | ✓ | ✓ | ✓ | | | | | | | | | |
| Lapatinib | ✓ | ✓ | ✓ | ✓ | ✓ | ✓ | ✓ | ✓ | ✓ | ✓ | ✓ | ✓ | ✓ | | | | | | | | | |
| Afatinib | ✓ | ✓ | ✓ | ✓ | ✓ | ✓ | ✓ | ✓ | ✓ | ✓ | ✓ | ✓ | ✓ | | | | | | | | | |
| Axitinib | ✓ | ✓ | ✓ | ✓ | ✓ | ✓ | ✓ | ✓ | ✓ | ✓ | ✓ | ✓ | ✓ | | | | | | | | | |
| Lestaurtinib | ✓ | ✓ | ✓ | ✓ | ✓ | ✓ | ✓ | ✓ | ✓ | ✓ | ✓ | ✓ | ✓ | | | | | | | | | |
| Nintedanib | ✓ | ✓ | ✓ | ✓ | ✓ | ✓ | ✓ | ✓ | ✓ | ✓ | ✓ | ✓ | | | | | | | | | | |
| Quercetin | | ✓ | | ✓ | | | | ✓ | | | | | | | | | | | | | | |
| Luteolin | | ✓ | | | | | | | | | | | | | | | | | | | | |
| Vemurafenib | ✓ | | | | | | | | | | | | | | | | | | | | | |
| Mebendazole | ✓ | | | | | | | | | | | | | | | | | | | | | |
| Neomycin | ✓ | | | | | | | | | | | | | | | | | | | | | |
| Triamterene | | | ✓ | | | | | | | | | | | | | | | | | | | |
| Mitoxantrone | | | | | | | | ✓ | | | | | | | | | | | | | | |
